# Gold-standard diagnostics are tarnished by lytic bacteriophage

**DOI:** 10.1101/2020.01.30.926832

**Authors:** E.J. Nelson, J.A. Grembi, D.L. Chao, J.R. Andrews, L. Alexandrova, P. H. Rodriguez, V.V. Ramachandran, Md.A. Sayeed, W. J. Wamala, A.K. Debes, D.A. Sack, A. J. Hryckowian, F. Haque, S. Khatun, M. Rahman, A. Chien, A.M. Spormann, G.K. Schoolnik

**Affiliations:** Departments of Pediatrics and Environmental and Global Health, University of Florida (USA); Department of Pediatrics, School of Medicine, Stanford University (USA); Department of Medicine, School of Medicine, Stanford University (USA); Department of Civil and Environmental Engineering, Stanford University (USA); Institute for Disease Modeling (USA); Vincent Coates Foundation Mass Spectrometry Laboratory, Stanford University (USA); Country Preparedness and IHR (CPI), World Health Organization (South Sudan); Johns Hopkins Bloomberg School of Public Health, Johns Hopkins University (USA); Department of Microbiology, School of Medicine, Stanford University (USA); Institute of Epidemiology, Disease Control and Research, Ministry of Health and Family Welfare, Government of Bangladesh. (Bangladesh); University College London (UK); International Centre for Diarrhoeal Disease Research, Bangladesh. (Bangladesh)

**Author notes:** Former institution (E.J. Nelson, J.A. Grembi, F. Haque, S. Khatun, M. Rahman). Address correspondence to Eric J. Nelson, MD PhD.

## Abstract

**Background:** A fundamental clinical and scientific concern is how lytic bacteriophage, as well as antibiotics, impact diagnostic positivity.

**Methods:** Cholera was chosen as a model disease to investigate this important question. Patients with diarrheal disease were enrolled at two remote hospitals in Bangladesh. Diagnostic performance was assessed as a function of lytic bacteriophage detection, as well as exposure to the first-line antibiotic azithromycin detected by mass spectrometry.

**Results:** Among diarrheal samples positive by nanoliter quantitative PCR for *Vibrio cholerae* (n=78/849), the odds that a rapid diagnostic test (RDT) or qPCR was positive was reduced by 89% (OR 0.108; 95%CI 0.002-0.872) and 87% (OR 0.130; 95%CI 0.022-0.649) when lytic bacteriophage were detected, respectively. The odds that a rapid diagnostic test (RDT) or qPCR was positive was reduced by more than 99% (OR 0.00; 95% CI: 0.00-0.28) and 89% (OR 0.11; 95% CI: 0.03-0.44) when azithromycin was detected, respectively.

**Conclusions:** Estimations of cholera burden may improve by accommodating for the negative effect of antimicrobial exposure on diagnostic positivity. Furthermore, the findings herein challenge our current approach to interpreting and developing bacterial diagnostics given variable rates of lytic bacteriophage and antibiotic exposure.

## BACKGROUND

There are approximately 4.5 billion diarrheal diseases cases per year [1]. While the 2-4 million cases of cholera that occur annually represent a small fraction of the total cases [2], cholera inflicts high morbidity and mortality on populations with extreme poverty. Outbreaks begin when immunologically susceptible human hosts are exposed to the Gram-negative pathogen *V. cholerae* (O1 and O139 serogroups) from contaminated food or water [3]. Before modern rehydration regimens, mortality rates rose above 20% [4] from acute secretory diarrhea resulting from the action of cholera toxin [3]. Treated with oral or intravenous rehydration, mortality rates decrease to less than one percent [5, 6]. Antibiotics are recommended for cholera patients with moderate to severe dehydration [7, 8], but in practice most cholera patients are likely ordered antibiotics. Asymptomatic cases are detected by a rise in antibody titer but negative stool studies [9]. Recovered patients become susceptible over time as a function of the durability of their immune response [3]. *V. cholerae* are shed from the human host with increased infectivity [10, 11]. This ‘hyper-infectivity’ is suggested to drive the exponential phase of outbreaks [12].

Patients can shed *V. cholerae* specific lytic bacteriophage (ICP1, 2, 3); ICP1 is specific for O1 *V. cholerae* [13, 14]. These vibriophages are proposed to quench outbreaks based on data that a higher percentage of patients shed vibriophage during the collapse of an outbreak [15–17]. Diagnostically, culture and PCR approaches are the best available ‘gold-standards’ for the detection of *V. cholerae* [18, 19]. Alternative methods include direct immuno-fluorescence microscopy for the O-antigen polysaccharide (OPS) [20], RDTs that rely on OPS specific antibodies, and recently by nl-qPCR [21, 22].

The rationale for this study was based on the recognition that cholera RDTs have limited adoption because of variable performance for unknown reasons [19, 23–26]; immediately testing stool samples demonstrated broad sensitivities (58-100%) and specificity (71-100%). A modified method enriches for *V. cholerae* in alkaline peptone water (APW) to increase specificity to 91-99% yet is associated with a decrease in sensitivity [23, 24, 27]. Both lytic phage and antibiotics have been postulated to impact diagnostics [26]. Using cholera as a model, we tested the hypothesis that lytic bacteriophage, and antibiotics, negatively impact diagnostics within the confines of a previously published clinical study [28]. In brief, the study was conducted from September to December 2015 at a district and sub-district hospital in the remote Northern district of Netrokona that is prone to seasonal cholera outbreaks. Inclusion criteria were patients at least two-months old and presented with acute (< 7 days) diarrhea (>3 loose stools in the 24 hours prior to admission) without complications.

## METHODS

### Subjects

This study was conducted with the confines of previously published studies in Bangladesh [28] and South Sudan [29]. Ethical approvals were obtained for the Bangladesh study at the Institutional Review Boards (IRBs) of Stanford University School of Medicine and the Institute of Epidemiology, Disease Control and Research, Bangladesh Ministry of Health and Family Welfare [28], and for South Sudan Study at the IRBs of Johns Hopkins Bloomberg School of Public Health and the South Sudan Ministry of Health, Directorate of Monitoring, Evaluation and Research [29]. Written informed consent was obtained from participants over 18 years, or guardians of participants.

### Clinical Study

In Bangladesh, inclusion criteria were patients at least two-months of age presenting with acute (< 7 days) diarrhea (>3 loose stools in the prior 24 hours) without clinical complications. Sample collection occurred from September to December 2015 at a district and sub-district hospital in the remote Northern district of Netrokona that is prone to seasonal cholera outbreaks. In South Sudan, inclusion criteria were patients presenting at a cholera treatment center in Juba who were at least 6 months-old, had diarrhea (>3 loose stools in the prior 24 hours) and no history of cholera vaccination. Samples were collected from August to September 2015.

### Laboratory Procedures

For samples collected in Bangladesh, the methods have been previously described [21, 22]. In brief, the first stool sample voided was collected immediately after admission to avoid exposure to hospital administered antibiotics. The supernatants from *V. cholerae* positive stools were tested for antibiotic exposure using a LC/MS protocol for a 1100 series HPLC (Agilent Technologies) integrated with an LTQ XL ion trap mass spectrometer (Thermo Fisher Scientific) [21]. The stool samples were tested by RDT (Crystal VC, Span Diagnostics) after enrichment in APW for 6 hours or overnight [28]. The first and last samples collected per day were stored in Cary-Blair media (4°C) for culture at a central reference laboratory in Dhaka (icddr,b); samples were stored for up to 1 month. Aliquots (500 μl) from all patients were stored in 1.3ml*RNAlater* (Invitrogen).

For Bangladesh samples, DNA was extracted using the MoBio Power Soil 96-well plate system (Qiagen; formerly PowerSoil). DNA extracts were screened in technical replicates for *V. cholerae* by qPCR in a 384-well Light Cycler (Roche) using *tcpA*^set1^ primers (Table S1) [21]. Samples that had CT values less than 25 were defined as positive. Samples with CT values from 25 to 31 were evaluated by PCR for *ompW* [8]. In parallel, nl-qPCR was performed in technical replicates with *tcpA*^set1^ and additional targets [21, 22]. Cyber Green master mix (Sigma Aldrich) was used for both qPCR and nl-qPCR however there was 1.8-fold more DNA in nl-qPCR reactions. Cycle thresholds for positivity for qPCR and nl-qPCR were 29 and 28, respectively. 16S rDNA analysis utilized previously published methods and data [21] on nl-qPCR *V. cholerae* positive samples for *tcpA* (Table S1). Lytic vibriophages ICP1, 2, and 3 were detected by PCR (Table S1). For samples collected in South Sudan, analyses for *V. cholerae* have been previously described on DNA extracted from dried stool spots [29]. In addition, the extracts were analyzed by PCR for ICP1 and ICP3 (ICP2 PCR technically failed; Table S1).

Direct immune-fluorescence was performed as previously described on planktonic cells from RNA*later* preserved stool samples [30]. This fraction was obtained by a 15 seconds 100-G centrifugation to remove sediment from 500 μl of sample, one PBS wash, pelleting the supernatant fraction, and resuspension of the pellet in 500 μl of PBS with 3.7% formalin. Mock positive control stool samples were used for molecular and microscopy assays that consisted of *V. cholerae* set to concentrations relative to cholera stool (5e8 CFU/ml and 1e8CFU/ml) in 500 ml normal saline plus 1.3 ml RNA*later*(ratio used in stool storage).

### Statistical analysis

Latent class modeling was used to estimate sensitivities and specificities of each diagnostic [31]. For prior information, the assumptions for sensitivities were the same for RDT, qPCR, nl-qPCR and culture (50-100%). Assumptions for specificities were 50-100% for RDT, 90-100% for qPCR and nl-qPCR, and 99-100% for culture [18]. Gibbs sampling with 100,000 iterations was used to generate posterior estimates with 95% credible intervals (CI). Fischer’s exact test was used to evaluate associations between diagnostic type and detection of lytic bacteriophage / azithromycin. Both sample odds ratios and estimated sample odds ratios with a conditional Maximum Likelihood Estimate were computed. A two-sample Wilcoxin test was used to compare CT values between diagnostic positive and negative samples among samples positive for *V. cholerae* positive by nl-qPCR CT. Comparison of microbiota (16S rDNA analysis) by diagnostic result and exposure among nl-qPCR positive samples was conducted by PERMANOVA as previously described [21]. Missingness in the dataset is designated as ‘NA’ and is restricted to laboratory results. Statistical analyses were completed in Graphpad Prism 8.0.1 and R v3.4.1 / RStudio v1.1.0153 [32].

### Data availability

Data analyzed in the manuscript have been made available in the online supplementary material.

## RESULTS

### Sensitivity and specificity estimates by latent class modeling

In Bangladesh, stool samples were collected from 881 of 961 enrolled patients. Among samples tested by RDT, qPCR, and nl-qPCR, the distribution of diagnostic positivity is provided (Fig 1A,B). The sensitivities and specificities of each diagnostic were estimated using a Bayesian latent class modeling framework, which enables estimation of diagnostic accuracy in the absence of a perfect reference standard by integrating data from multiple tests [31]. Estimates for sensitivity of RDT, qPCR, and nl-qPCR were 31.5% (95% CI:21.5–43.7), 64.1% (CI: 50.7-80.2) and 97.6% (95% CI: 89.0–100.0), respectively. The specificities were 99.6% (95% CI: 99.0–99.9), 99.9% (95% CI: 99.7–100.0) and 99.6% (95% CI: 98.3–100.0), respectively. Among the subset of samples randomly chose for culture (16 positive out of 251), sensitivity was 57.1% (40.4-73.2) and specificity 99.7 (99.3-99.9). Based on these results, nl-qPCR was selected as the best available reference standard for subsequent analysis and the receiver operator curve (ROC) is presented (Fig 1C).

**Figure 1.**
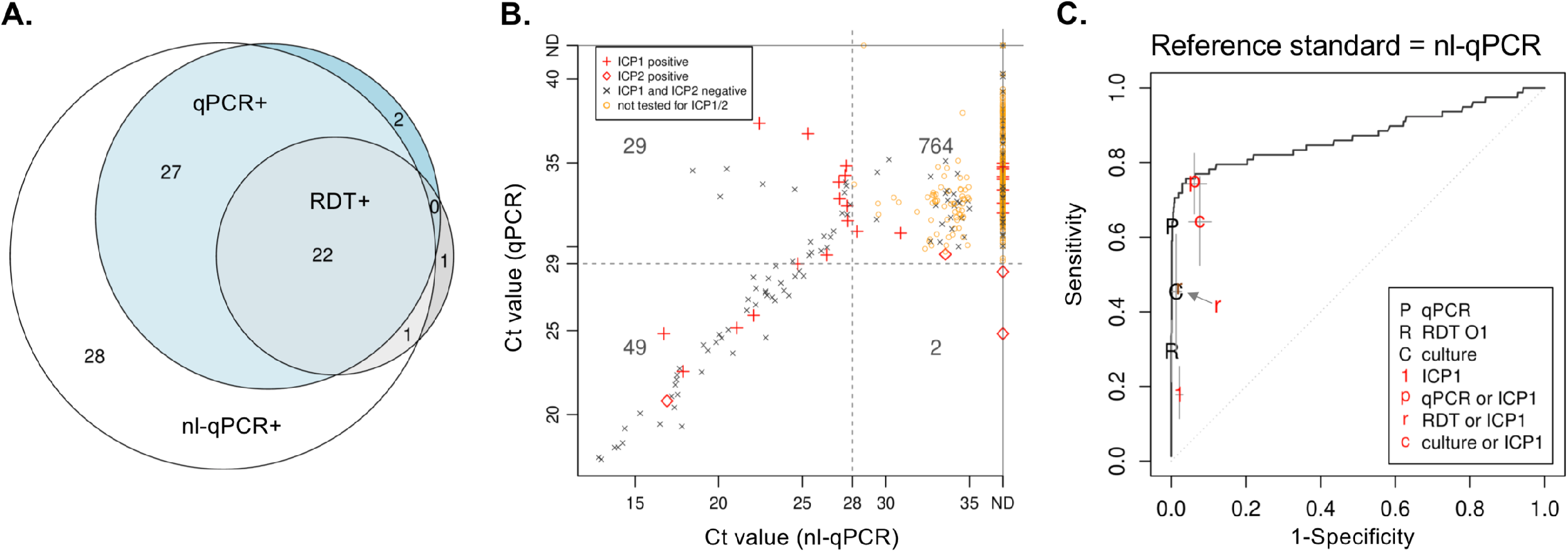
Diagnostic evaluation. **A.** Euler diagram of diagnostic positivity for qPCR, nl-qPCR, and RDT; area within each circle is relative to the degree of positivity. **B.** Comparison Ct values between qPCR and nl-qPCR analysis with ICP1 and ICP2 metadata; horizontal and vertical dotted lines depict thresholds of positivity for each test; ND= not detected. **C.** Receiver operator characteristic (ROC) curve. Estimates of the sensitivity and 1-specificity of combining diagnostics are defined in the key and vertical bars from each symbol depict the 95% CI.

### Impact of lytic phage on diagnostic positivity

Among *V. cholerae* positive samples by nl-qPCR, 19.2% (15/78) and 1.3% (1/78) were positive for ICP1 and 2, respectively; ICP3 was not detected. Of 180 random samples negative by nl-qPCR, qPCR and RDT, two patients had ICP1 (one was culture positive) and one had ICP2. Among *V. cholerae* positive samples by nl-qPCR that lacked azithromycin, vibriophage were negatively associated with diagnostic positivity by RDT (OR 0.11; 95% CI: 0.002-0.87), qPCR (OR 0.13; 95% CI: 0.02-0.65) and direct immuno-fluorescent microscopy [30] (OR 0.18; 95% CI 0.02-1.031; Table 1). Frequencies of vibriophage detection were different between study sites (Fischer’s exact test; p = 0.033). Diarrheal samples from South Sudan were analyzed to increase generalizability [29]. ICP1 was detected in 10.2%(n=10/98) of all enriched samples, 24% of samples (n=7/29) that were PCR positive samples for *V. cholerae* and 5.7% (n=3/69) of samples that were RDT negative by PCR for *V. cholerae*. ICP1 was negatively associated with RDT positivity after enrichment (OR 0.00, 95%CI 0.00-0.64, p=0.010; Table S2); a statistically significant difference was not observed for unenriched samples. ICP3 was not identified. There were insufficient samples to assess phage impact on culture positivity.

**Table 1.**
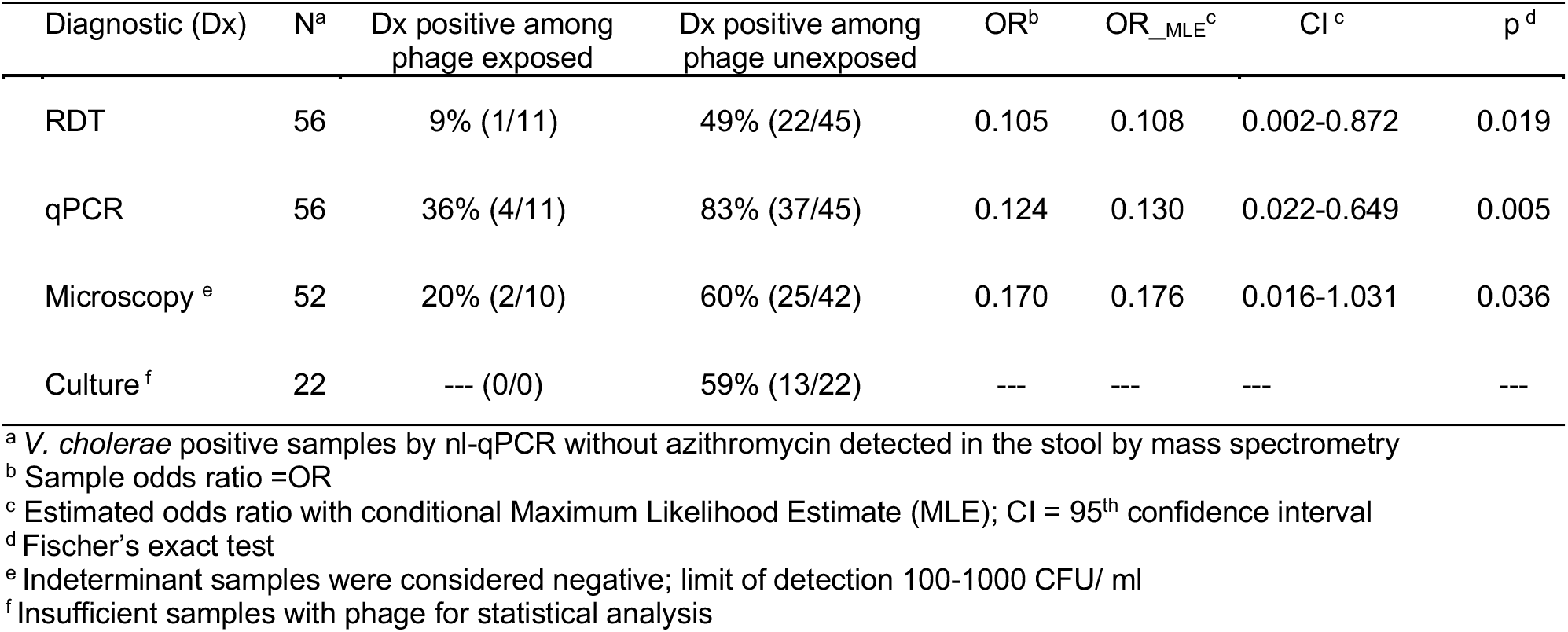
Lytic phage negatively impact diagnostic positivity (azithromycin excluded)

### Impact of azithromycin on diagnostic positivity

Among Bangladesh samples positive by nl-qPCR but negative for bacteriophage, azithromycin was negatively associated with diagnostic positivity by RDT (OR 0.00; 95% CI: 0.00-0.28) and qPCR (OR 0.11; 95% CI: 0.03-0.44), but not by direct fluorescent microscopy (OR 0.54; 95% CI 0.14-1.97; Table 2). Azithromycin was negatively associated with culture positivity (OR 0.00, 95% 0.00-0.997; Table 2).

**Table 2.** Azithromycin negatively impacts diagnostic positivity (phage excluded)

| Diagnostic (Dx) | N<sup>a</sup> | Dx positive among azithro exposed | Dx positive among azithro unexposed | OR<sup>b</sup> | OR_MLE<sup>c</sup> | CI<sup>c</sup> | p<sup>d</sup> |
| --- | --- | --- | --- | --- | --- | --- | --- |
| RDT | 63 | 0% (0/18) | 49% (22/45) | 0.000 | 0.000 | 0.000-0.282 | <0.001 |
| qPCR | 63 | 33% (6/18) | 82% (37/45) | 0.108 | 0.113 | 0.026-0.437 | <0.001 |
| Microscopy<sup>e</sup> | 58 | 44% (7/16) | 60% (25/42) | 0.529 | 0.535 | 0.139-1.973 | 0.378 |
| Culture<sup>f</sup> | 27 | 0% (0/5) | 59% (13/22) | 0.000 | 0.000 | 0.000-0.997 | 0.041 |

<sup>a</sup> *V. cholerae* positive samples by nl-qPCR without lytic bacteriophage (ICP1,2,3) detected in the stool by PCR
<sup>b</sup> Sample odds ratio =OR
<sup>c</sup> Estimated odds ratio with conditional Maximum Likelihood Estimate (MLE); CI = 95<sup>th</sup> confidence interval
<sup>d</sup> Fischer's exact test
<sup>e</sup> Indeterminant samples were considered negative; limit of detection 100-1000 CFU/ ml
<sup>f</sup> Insufficient samples with phage for statistical analysis

### Absolute and relative *V. cholerae* concentration

Absolute and relative *V. cholerae* concentration was assessed by nl-qPCR and 16S rDNA analysis, respectively. Among nl-qPCR positive samples, there was a significant inverse relationship between diagnostic positivity and *V. cholerae* concentration (Table S3). With no exclusions, fold-differences between positive and negative samples ranged from 21-fold (culture) to 79-fold (qPCR). The one exception was that phage exposure (azithromycin samples excluded) did not associate with a significant difference in the nl-qPCR Ct values between culture positive (n=13; Ct = 19.4, 95% CI 14.3-22.0) and negative samples (n=9; Ct =20.8, 95% CI 17.6-25.9; p=0.186). Statistically significant differences in microbiota (16S rDNA) were observed between RDT positive and negative stools with stratifications for bacteriophage (Fig S2A) and azithromycin (Fig S2B).

## DISCUSSION

This study investigated the potential vulnerability diagnostics have when bacterial targets are exposed to lytic bacteriophage predation or antibiotics. Using cholera as a model system and nl-qPCR as a reference standard for *V. cholerae,* we found that the odds of a RDT, qPCR and microscopy diagnostic testing positive were reduced by more than 83% when lytic bacteriophage were present. Similarly, the odds of a RDT, qPCR and culture testing positive were reduced by more than 89% when the first-line antibiotic azithromycin was detected in stool by mass spectrometry. These results expose a vulnerability of gold-standard diagnostics that clinicians and microbiologists feared but lacked sufficient data to take evidence-based action.

We reason that the low inflection point in the ROC at approximately 0.7 sensitivity is multi-factorial (Fig 1C). We explored the effect on sensitivity and specificity of adding ICP1 detection as a proxy for *V. cholerae* detection (Fig 1C). Both qPCR, culture and the RDT moderately improved. The effects of lytic bacteriophage, antibiotics and host antimicrobial factors on diagnostic positivity are likely additive, especially given that these diagnostics target different biologic mechanisms. How duration of illness and severity of disease serve as determinants of diagnostic positivity also remain unknown. Time-series analyses of cholera patients with defined antimicrobial exposures are needed steps to further these lines of inquiry.

These findings should be viewed within the context of the limitations of the study. The procedures were chosen for feasibility at remote field sites. This delayed cultures up to one-month and precluded plaque assays. The higher detection rate of nl-qPCR compared to qPCR was multi-factorial, including the 1.8-fold difference in DNA. The positive nl-qPCR samples that were negative by qPCR and negative by *ompW* were unlikely to be false positives because *Vibrio* spp. were detected by 16S rDNA analysis in all 13 samples that did not have lytic vibriophage; those with vibriophage did not result (n=7/7). Among nl-qPCR positive and qPCR negative samples, PCR detection for *tpcA* correlated with PCR detection of *ctxA* (cholera toxin;n=5/5; Table S1). These toxin data, paired with serologic results that found only O1 *V. cholerae*, makes the possibility of confounding from non-O1 *V. cholerae* unlikely. Despite these limitations, the discovery that lytic bacteriophage negatively impacts diagnostics, even to the point that samples will test positive for bacteriophage and negative for the pathogen, has broad significance. One explanation is lytic bacteriophage and antibiotics inhibit bacterial growth below the diagnostic limits of detection. Alternatively, bacteriophage nucleases, or host nucleases responding to bacteriophage infection, may digest host chromosomal DNA to the point that PCR fails [33, 34].

### Conclusion

Within the cholera field, this study suggests that more nuanced analytical approaches are needed to determine the true cholera burden during outbreaks, especially in the latter phases when rates of concurrent lytic bacteriophage predation are likely higher [16, 17]. This may require an approach that includes lytic bacteriophage detection as a proxy for pathogen detection and a de-emphasis on diagnostic results with known antibiotic exposure. Outside the cholera field, these data serve as a call-to-action to survey for lytic bacteriophage when bacterial diagnostics have inconsistent performance, especially when there is discordance between clinical presentation and diagnostic result. These efforts may justify a new line of diagnostic development that targets both the prey (pathogen) and predator (bacteriophage).

## Acknowledgements

We thank the patients for participating in this study, the field and central project staff, and diligent clinical team who made this study possible. We are grateful to M. Barry, S. Luby and Y. Maldonado for their support at Stanford University, as well as G. Morris, A. Ali, M. Alam, E. Schmidt, R. Autrey, K. Berquist and S. Rivkees at the University of Florida for their support, reagents and helpful discussions.

## Disclaimer

These funders had no role in study design, data collection and analysis, decision to publish, or preparation of the manuscript.

## Supplement Material

**Figure S1.**
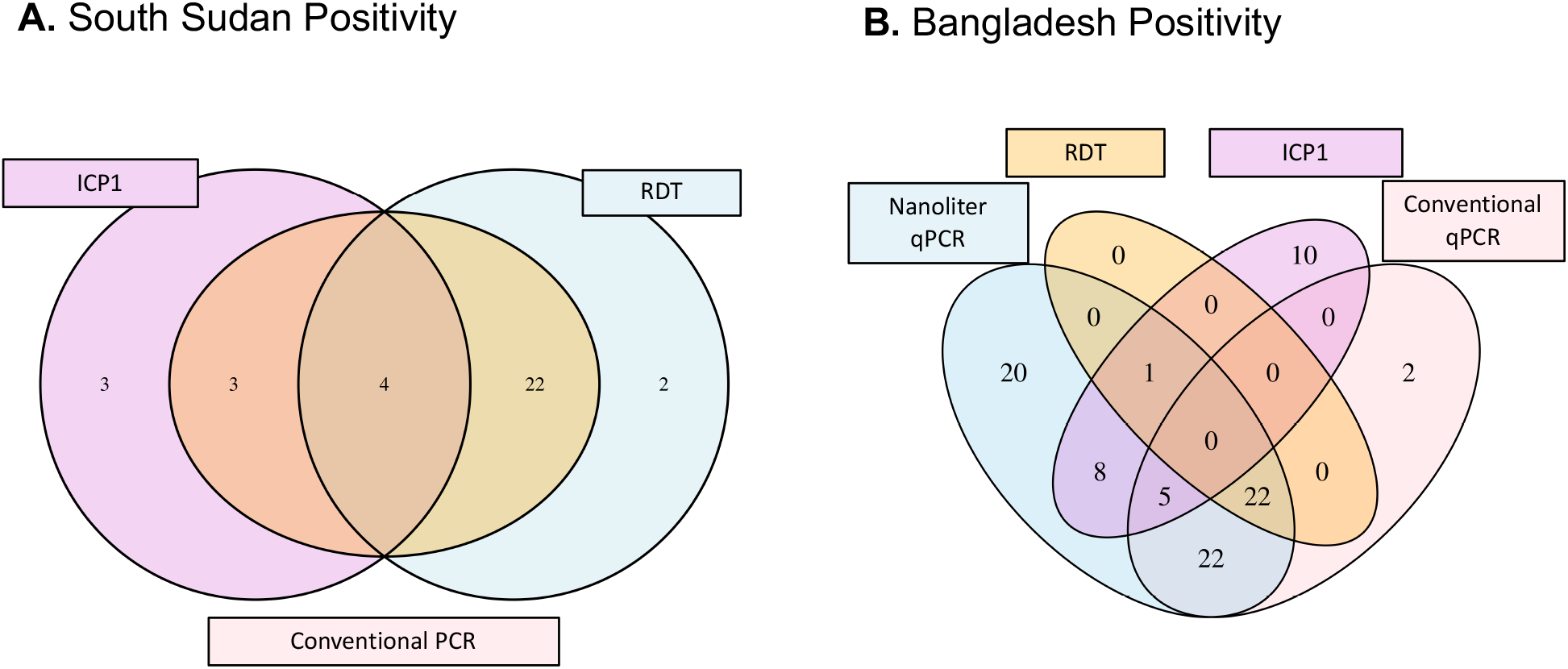
Comparison of diagnostic positivity and ICP1 detection in the libraries from South Sudan **(A)** and Bangladesh **(B)**. RDT= rapid diagnostic test. Both settings used the Crystal VC test with enrichment. Data presented from S. Sudan is based on PCR performed at Institute Pasteur (Table S2).

**Figure S2.**
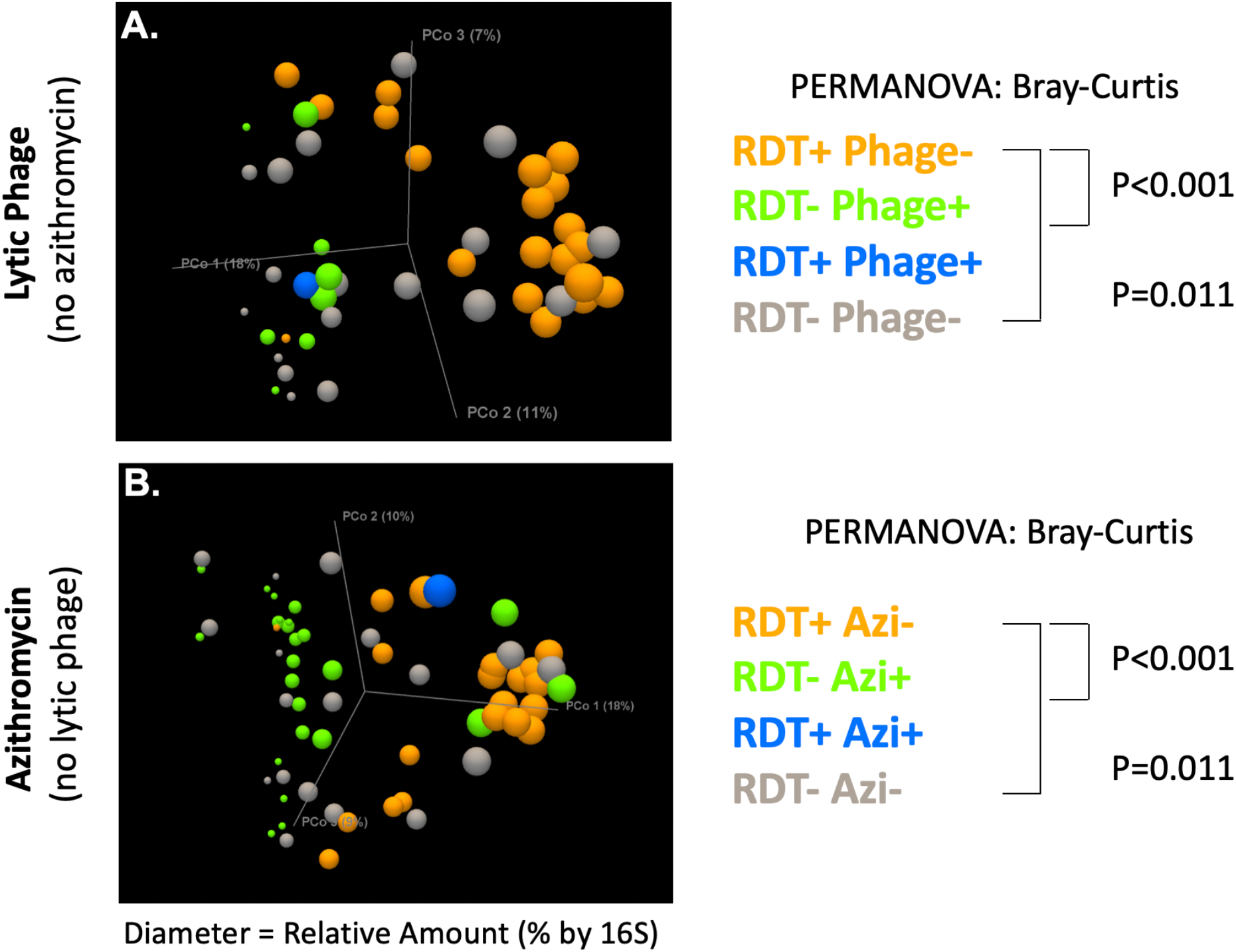
Principal component analysis of 16S rDNA analysis of *V. cholerae* positive samples by nl-qPCR analysis as previously described[21]. **A.** Among samples without azithromycin detection, data are color coded based on RDT and phage positivity (right). **B.** Among samples without bacteriophage detection, data are color coded based on RDT and azithromycin positivity (right). PC1 is oriented horizontally and the icon size is set to the relative percentage of *V. cholerae* detected in the microbiota (0-25%, 26-50%, 51-75%, 76-100%). For both the upper and lower panels, statistically significant differences between groups were detected by PERMANOVA (Bray-Curtis)[21]. *V. cholerae* positivity is defined by nl-qPCR positivity with either tcpA primer sets to be consistent with prior analytic approach[21].

**Table S1.**
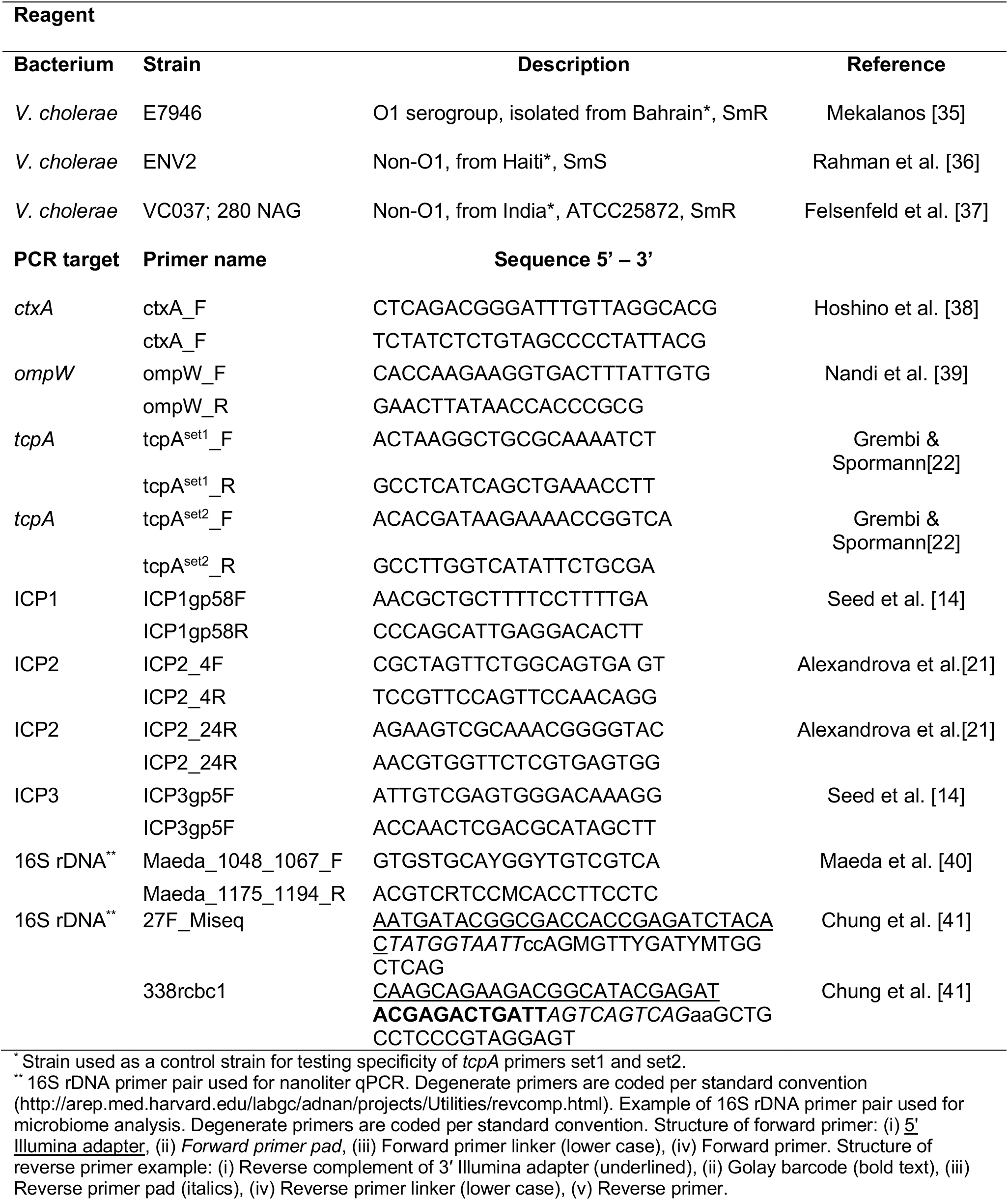
Reagents

**Table S2.** South Sudan: Impact of bacteriophage ICP1 on cholera RDT positivity

| Diagnostic Test | Total | Diagnostic Positive |  | Diagnostic Negative |  | OR<sub>MLE</sub><sup>a</sup> | CI<sup>a</sup> | P<sup>b</sup> |
| --- | --- | --- | --- | --- | --- | --- | --- | --- |
|  |  | Phage Pos | Phage Neg | Phage Pos | Phage Neg |  |  |  |
| Among VC PCR positive (Hopkins)<sup>c</sup> |  |  |  |  |  |  |  |  |
| RDT (Direct)<sup>d</sup> | 32 | 7 | 26 | 1 | 0 | --- | --- | --- |
| RDT (Enriched) | 34 | 5 | 25 | 3 | 1 | 0.075 | 0.001-1.137 | 0.033 |
| Among VC PCR positive (Pasteur)<sup>c</sup> |  |  |  |  |  |  |  |  |
| RDT (Direct)<sup>d</sup> | 29 | 6 | 22 | 1 | 0 | --- | --- | --- |
| RDT (Enriched) | 29 | 4 | 22 | 3 | 0 | 0.000 | 0.000-0.641 | 0.010 |

<sup>a</sup> Estimated odds ratio with conditional Maximum Likelihood Estimate (MLE); CI = 95<sup>th</sup> confidence interval
<sup>b</sup> Fischer's exact test
<sup>c</sup> Conventional PCR was performed by Johns Hopkins University and Institute Pasteur for cross validation [29].
'Direct' = tested directly from stool; 'Enriched' = grown in selective media (APW) prior to testing. Values reported in
the main text and Figure S1 are derived from the Institute Pasteur data set.
<sup>d</sup> Insufficient sample size for statistical analysis

**Table S3.** Fold-differences in target DNA detection among *V. cholerae* positive samples

|  |  | Diagnostic Positive | Diagnostic Negative |  |  |
| --- | --- | --- | --- | --- | --- |
| Diagnostic Test | Total<sup>1</sup> | Median nl-qPCR Ct (n; IQR) | Median nl-qPCR Ct (n; IQR) | Fold-difference | p<sup>2</sup> |
| No exclusion |  |  |  |  |  |
| RDT | 78 | 19.0 (23; 16.5-21.7) | 24.4 (55; 21.1-27.7) | 42 | <0.001 |
| qPCR | 78 | 20.2 (49; 17.4-22.9) | 26.5 (29; 24.9-27.6) | 79 | <0.001 |
| Microscopy<sup>3</sup> | 71 | 20.3 (34; 17.3-23.0) | 25.1 (37; 21.5-27.3) | 28 | <0.001 |
| Culture | 29 | 19.4 (13; 14.3-22.0) | 23.8 (16; 17.7-25.5) | 21 | 0.013 |
| Includes samples with azithromycin (excludes phage samples) |  |  |  |  |  |
| RDT | 63 | 18.4 (22; 16.2-21.1) | 23.9 (41; 21.3-26.2) | 46 | <0.001 |
| qPCR | 63 | 20.2 (43; 17.4-22.9) | 26.2 (20; 24.5-27.5) | 65 | <0.001 |
| Microscopy<sup>3</sup> | 58 | 20.3 (32; 17.4-23.2) | 24.5 (26; 20.7-26.2) | 18 | 0.002 |
| Culture | 27 | 19.4 (13; 14.3-22.0) | 23.8 (14; 17.8-25.5) | 21 | 0.011 |
| Includes samples with phage (excludes azithromycin samples) |  |  |  |  |  |
| RDT | 56 | 19.0 (23; 16.5-21.7) | 24.6 (33; 21.1-27.0) | 48 | <0.001 |
| qPCR | 56 | 19.9 (41; 17.4-22.8) | 27.2 (15; 25.4-27.6) | 163 | <0.001 |
| Microscopy<sup>3</sup> | 52 | 19.1 (27; 17.2-22.8) | 24.8 (25; 21.5-27.2) | 53 | <0.001 |
| Culture | 22 | 19.4 (13; 14.3-22.0) | 20.8 (09; 17.6-25.9) | 2.8 | 0.186 |

<sup>1</sup> Nanoliter (nL) qPCR positive for *V. cholerae* (Ct < 28); a random subset were cultured or available for microscopy.
<sup>2</sup> Mann-Whitney U test (two-tailed)
<sup>3</sup> Indeterminant samples scored as negative; limit of detection 100-1000 CFU/ ml

